# Mitochondrial and metabolic remodeling in human skin fibroblasts in response to glucose availability

**DOI:** 10.1101/2021.02.24.432508

**Authors:** Cláudio F. Costa, Sónia A. Pinho, Sonia L.C. Pinho, Inês Miranda-Santos, Olivia Bagshaw, Jeffrey Stuart, Paulo J. Oliveira, Teresa Cunha-Oliveira

## Abstract

Cell culture conditions highly influence cell metabolism *in vitro*. This is relevant for preclinical assays, for which fibroblasts are an interesting cell model, with applications in regenerative medicine, diagnostics and therapeutic development for personalized medicine as well as in the validation of ingredients for cosmetics. Given these cells’ short lifespan in culture, we aimed to identify the best cell culture conditions and promising markers to study mitochondrial health and stress in Normal Human Dermal Fibroblasts (NHDF). We tested the effect of reducing glucose concentration in the cell medium from high glucose (HGm) to a more physiological level (LGm), or its complete removal and replacement by galactose (OXPHOSm), always in the presence of glutamine and pyruvate. We have demonstrated that only with OXPHOSm it was possible to observe the selective inhibition of mitochondrial ATP production. This reliance on mitochondrial ATP was accompanied by changes in oxygen consumption rate (OCR) and extracellular acidification rate (ECAR), oxidation of citric acid cycle substrates, fatty acids, lactate and other substrates, mitochondrial network extension and polarization and changes in several key transcripts related to energy metabolism. We also evaluated the relevance of galactose, glutamine and pyruvate for OXPHOS stimulation, by comparing OCR and ECAR in the presence or absence of these substrates. Galactose and pyruvate seem to be important, but redundant, to promote OXPHOS, whereas glutamine was essential. We concluded that LGm does not promote significant metabolic changes but the short-term adaptation to OXPHOSm is ideal for studying mitochondrial health and stress in NHDF.

**Author Contributions:** CC, SAP, SLCP and IMS performed experiments. TCO and PJO designed research and acquired funding. JS, and OB analyzed data. CC and TCO analyzed data and wrote the paper. All authors contributed to the final version of the manuscript.

## Introduction

Cell culture plays an essential part in preclinical trials and actively contributes to reducing the use of animals in research. However, it has recently become apparent that traditional commercial media for human cell culture does not accurately replicate extracellular fluid in vivo affecting many aspects of cell biology^1^. This has important implications for cancer biology. However, for other cell culture applications, experimentation may benefit from focusing on other aspects of media compositions. For example, the study of mitochondrial diseases, toxicity, and development of mitochondria-targeted drugs may benefit more from media that promote cellular respiration.

Isolated human skin fibroblasts are routinely used to characterize mitochondrial diseases and as experimental models for drug development. These cells can be easily obtained from patients, including those with mitochondrial disorders, by minimally invasive methods^2^. Fibroblasts from patients with mitochondrial deficiencies can be a good model for characterizing the associated metabolic disturbance and developing therapeutic approaches for personalized medicine^2, 3, 4^. Normal human skin fibroblasts may also be an interesting toxicological model for testing ingredients for cosmetics^5, 6^, which is an important application of mitochondria-targeted compounds (see patents # US10085966B2, US9427444B2, WO2018069904A1 or WO2018122789A1 for examples). However, their penchant in the cell culture environment for glucose fermentation over complete oxidation can make them less susceptible to respiration deficits. Stimulation of OXPHOS in cultured cells can be achieved by replacing glucose with galactose in the culture medium. This approach has been used to reveal mitochondrial toxicity^7^, including that underlying drug-induced liver injury^8^. However, this strategy does not work for all cell lines. C2C12 mouse myoblasts, for example, do not grow well in galactose-based DMEM in the absence of glutamine. Instead, increased reliance on OXPHOS can be achieved by maintaining glucose in the culture medium at physiological levels of 5 mM (low glucose; LG)^9^. Using either strategy, adopting a more oxidative phenotype in cells with endogenous or exogenous mitochondrial deficiencies can highlight the deleterious consequences of these deficiencies. Hence, this approach is useful for diagnosing, characterizing, and developing treatments for mitochondrial deficiencies and toxicities^10, 11^. Commercial primary human fibroblast cell lines can grow in OXPHOSm^12^. However, primary skin fibroblasts have a short life-span in culture and protocols that involve adaptation to media containing galactose are usually long, taking most of these cells’ available lifespan^12^. Thus, it is essential to develop fast, robust, and reproducible screening assays that rapidly ‘rewire’ cultured primary human dermal fibroblasts to promote mitochondrial respiration. Here, we characterize two protocols for promoting reliance on oxidative metabolism in human dermal fibroblasts: acclimation to either a low-glucose, or a glucose-free galactose-containing DMEM. We determined if the absence of glucose induces mitochondrial remodeling and cellular reliance on mitochondria for ATP production to assess if the reduction of glucose availability to physiological levels is enough to promote mitochondrial remodeling and activity and if the removal and the reduction of glucose have comparable mitochondrial effects. We also determined how different energy substrates present in the culture medium impacted NHDF metabolic remodeling.

## Material and Methods

### Cell Culture

NHDF cell line (#CC-2511) was purchased from Lonza (Basel, Switzerland). NHDF cells were cultured at 37°C in a humidified atmosphere of 5% CO_2_ in High-Glucose Medium (HGm; Dulbecco’s Modified Eagle’s Medium (DMEM D5030) supplemented with the sugar 4.5 g/L (25 mM) D-glucose, 0.976 g/L (6 mM) L-glutamine, 0.11 g/L (1 mM) sodium pyruvate, 3.7 g/L (44 mM) sodium bicarbonate, 100 U/ml penicillin, 100 μg/ml streptomycin, 250 ng/ml antifungal amphotericin B and 10% (v/v) fetal bovine serum (FBS), pH 7.2). All cells were passaged by trypsinization at about 90% confluence.

Part of the cells was gradually adapted to the new media during one cell passage (**Supplemental Figure 1**). Cells were plated in a mixture of HGm and final medium (OXPHOSm or Low-Glucose Medium (LGm) which are similar to HGm, but, instead of 25 mM glucose these media were supplemented with 1.8016 g/L (10 mM) galactose or 0.9 g/L (5 mM) D-glucose, respectively), in a ratio of 75:25, in 100 mm-diameter tissue culture dishes at 37°C in a humidified atmosphere of 5% CO_2_. When about 50-60% confluence was reached, culture medium was replaced, without trypsinization, to a mixture of HGm and final medium, in a ratio of 50:50. When cells reached about 70-75% of confluence, culture medium was replaced, without trypsinization, to a mixture of HGm and final medium, in a ratio of 25:75. Finally, when about 90% of confluence was achieved, cells were passaged by trypsinization to 100% final medium. As recommended by the supplier, cells were used for various assays until a maximum of 15 passages from the initial vial.

For some experiments, cells were also adapted to culture media similar to OXPHOSm, but without galactose (-Gal), glutamine (-Gln), pyruvate (-Pyr), or galactose and pyruvate (-Gal – Pyr) supplementation.

### Analysis of bioenergetics using Seahorse MitoStress test

Oxygen consumption rate (OCR) and extracellular acidification rate (ECAR) were determined using a Seahorse XFe96 analyzer and the Seahorse XF Cell Mito Stress Test (Agilent Technologies, Santa Clara, CA). Cells were seeded into Seahorse XFe96 well plates at a density of 7500 cells/well/80 μl. One hour prior to the assay the plate was rinsed twice with non-supplemented DMEM D5030, 175 μl of the same culture medium was added and cells were incubated for 1 h at 37 °C without CO_2_. The corresponding culture medium was used in the assay, including in the preparation of the loaded compounds. A constant volume of 25 μl of each compound was pre-loaded into the respective ports of the cartridge (Port A: oligomycin, final concentration: 1 μM; Port B: FCCP, final concentration: 4 μM; Port C: a mix of rotenone/antimycin A, final concentration: 2 μM). Measurements were performed in five to six replicates. OCR and ECAR values were normalized to nuclei number and analyzed using Wave v2.3 software (Agilent Technologies, Santa Clara, CA). At the end of the experiment, cells were fixed with 10% (w/v) TCA, labeled with bisbenzimide trihydrochloride (Hoechst 33342) at 0.5 μg/mL, imaged in an InCell Analyzer 2200 (GE Healthcare, Chicago, Illinois, USA) and counted using IN Cell Analyzer 1000 analysis software.

### Intracellular ROS levels

General intracellular oxidative stress was determined by following the oxidation of the CM-H_2_DCFDA fluorescent dye (Molecular Probes, Invitrogen). NHDF cells grown in HGm, LGm and OXPHOSm were plated in 96-well black plates with optical bottoms at a cellular density of 3750 cells/well (18750 cells/mL). After 48 h, cells were incubated for 30 minutes at 37°C with culture medium supplemented with CM-H_2_DCFDA 5 μM but without sodium bicarbonate nor FBS. After CM-H_2_DCFDA loading, medium was replaced by new one without dye, sodium bicarbonate or FBS, and the kinetics of the fluorescence signal was recorded in a BioTek® Cytation3 UV–vis multi-well plate imaging reader (BioTek, Winooski, Vermont, USA) for 1 h, with excitation and emission wavelengths of 485 nm and 528 nm, respectively. Five independent experiments were done, each one with 6-8 technical replicates. At the end of the kinetic readings, cells were fixed with TCA 10%, nuclei were labelled with Hoechst 33342 at 0.5 μg/mL, imaged in an InCell Analyzer 2200 (GE Healthcare, Chicago, Illinois, USA) and counted using IN Cell Analyzer 1000 analysis software, Developer Toolbox. For each well, we calculated the difference between the fluorescence readings after 1 h (F) and the initial reading (F0) and divided per nuclei number, as in the equation (F-F0)/cell count.

### Metabolic Microarrays

#### Mito S1-Analysis of mitochondrial metabolic activity and substrate preference

To investigate mitochondrial metabolic activity and substrate preference we used the Mitoplate S-1 technology from Biolog, Inc. (Biolog#14105), consisting in 96-well plates preloaded with triplicates of 31 different substrates including TCA cycle intermediates, hexoses, trioses, amino acids and fatty acids. The Mitoplate S-1 assay assesses the electron flow rate through the electron respiratory chain resulting from the metabolization of each substrate pre-loaded in the respective well. It is a colorimetric assay based on the reduction of the tetrazolium redox dye MC, which acts as the final electron acceptor of the respiratory chain.

The assay-mix was prepared at twice the final concentration, consisting in Biolog MAS 2x (Biolog#72303), redox dye MC 6x (Biolog#74353), saponin 24x (Sigma #SAE0073, at 1.2 g/mL; final concentration of 50 μg/mL, for cell permeabilization) and sterile milliQ H_2_O. Mitoplates S-1 were activated by adding 30 μL of assay-mix 2x per well and incubated at 37° C, 5% CO_2_ for 1-2 h. For this assay, cells were trypsinized washed in PBS and resuspended in 1 mL Biolog MAS (1x) at 1 × 10^6^ cells/mL. Cells were plated at a density of 30 × 10^3^ cells/well from an inoculation volume of 30 μL/well. The number of cells per plate was previously optimized in order to have both good signal and not overload in all the wells. Dye MC reduction was followed kinetically for 24 h, with 5 min intervals, using the microplate reader OmniLog (Biolog). The initial rate was calculated in the linear phase (first hour) using the Biolog software data analysis® 1.7.

#### Mito I1-Analysis of cellular sensitivity to mitochondrial inhibitors

To investigate cellular sensitivity to mitochondrial inhibitors, we used the Mitoplate I-1 technology from Biolog, Inc. (Biolog#14104), consisting in 96-well plates pre-loaded with 22 mitochondrial inhibitors, each at 4 different concentrations, and the respective negative and positive control in quadruplicate. The Mitoplate I-1 assay assesses the rate of electron flow through the electron respiratory chain of energized mitochondria in the presence of specific inhibitors.

Assay-mix 2x was prepared as for Mitoplates S1, but including sodium succinate 24x (Fisher Scientific, S/6480/53, at 96 mM, final concentration of 2 mM), for mitochondrial energization. Mitoplates I-1 were activated by adding 30 μL of this assay-mix 2x per well. Cells were then prepared and added as described for Mitoplates S-1. Dye MC reduction was followed as described for Mitoplates S1, and the percentage of inhibition in each well was calculated using the dye reduction rates and following the equation: 100 x (well-negative control)/(positive control).

#### Fluorescence microscopy imaging

NHDF cells were plated at a density of 2000 cells/well in 96-well plates. Twenty four hours after plating, cells were incubated with 100 nM Tetramethylrhodamine Methyl Ester (TMRM) and 0.5 μg/ml Hoechst 33342 in medium without FBS, for 15 min in the dark, at 37°C and with a 5% CO_2_ atmosphere. Mitochondrial network and nuclei number were then assessed using InCell Analyzer 2200 (GE Healthcare, Chicago, Illinois, USA). Images were acquired in 16 fields per well at 10x magnification (Nikon 10x/0.45, PlanApo, CFI/60) in combination with two fluorescence detection channels (DAPI and CY3). Image processing was performed using ImageJ 1.51u software and the images were analyzed to obtain nuclei number counting and TMRM fluorescence, as an indirect measurement of the mitochondrial membrane potential, using the IN Cell Analyzer 1000 analysis software, Developer Toolbox.

#### Mitochondrial Network Analysis

For mitochondrial network morphology analysis, NHDF cells grown in HGm, LGm and OXPHOSm were plated in IBIDI μ-Slide 8 Well ibiTreat at a density of 6000 cells/well and after 24 h cells were incubated for 90 minutes with MitoTracker Red CMXRos (Molecular Probes, Invitrogen) at 100 nM, 37°C in the CO_2_ incubator. At the end, the probe was removed, and cells were fixed with 4 % paraformaldehyde and labelled with Hoechst 33342 at 0.5 μg/mL for nuclei detection.

Images were acquired on a Zeiss LSM 710 confocal microscope (Carl Zeiss, Jena, Germany) using a 63x magnification (Zeiss 63x/1.4, PlanApo DIC M27). MitoTracker Red CMXRos and Hoechst 33342 were detected using the 561 nm (DPSS 561-10 laser and 405 nm Diode 405-30 laser, respectively. The fluorescent light source was an HXP 120V. The pinhole aperture was set to 1 AU. Z-stacks were processed to obtain maximum z-projections and were acquired using the Zen Black 2012 software (Zeiss) with a step size of 0.41 – 0.46 μm.

Mitochondrial network morphology was analyzed using the Mitochondrial Network Analysis (MiNA) tool for the Fiji distribution of ImageJ^13^. A pre-processing macro was applied to the images to improve contrast of mitochondrial structures and reduce background signal. The contrast of mitochondrial structures was enhanced by applying the Tubeness filter (sigma=3 pixels)^14^. The signal intensity was then normalized to a common minimum and maximum before processing with MiNA. Briefly, images underwent thresholding using Li’s method to produce a binary image^15^. A morphological skeleton was then produced using the Skeletonize 2D/3D plugin^16, 17^. This method uses iterative thinning to remove outer pixels, resulting in a skeleton of mitochondrial structures, one pixel wide. Length measurements were extracted from the morphological skeleton using the Analyze Skeleton plugin and used to compute morphologically relevant parameters^17^. Mitochondrial footprint was quantified from the total area of mitochondrial signal-positive pixels. Mitochondria form branching networks where branches are separated by a node. Multiple independent networks; networks not attached by a branch, are found within one cell. All mitochondrial structures deduced from the morphological skeleton were used to calculate mean branch length and mean summed branch length per individual structure. Mean branch length was calculated as the average length of mitochondrial structures between two nodes. The mean sum of branch lengths per individual network was calculated by computing the sum of lengths between two nodes within an independent structure/network and dividing this by the total number of independent structures. This analysis was performed on a per cell basis. For each condition, 105-127 cells were analyzed.

### Quantification of Adenine nucleotides levels

#### Assessment of ATP/ADP and energy charge

For the initial characterization of cellular metabolism in different media, measurement of adenine nucleotide levels was performed by high performance liquid chromatography (HPLC). NHDF were grown in 100 mm plates for nucleotide extraction until about 80% confluence and washed with PBS 1x. Then, 1 ml of ice-cold 0.6 M HClO_4_ was added to each dish and content (lysates) was scraped to 2 ml tubes. All the procedure was performed on ice or at 4°C. Lysates were then centrifuged for 10 min at 14000 g, at 4°C. The resulting pellet was resuspended in 500 μl of 1 M NaOH for protein quantification, while the supernatant was neutralized with 3 M KOH/1.5 M Tris, and centrifuged again for 10 min at 14000 g, at 4°C. The resulting pellet was discarded and the supernatant was collected and immediately analyzed by reverse-phase HPLC, using a LiChrospher® 100 RP-18 (5 μm) LiChroCART® 125-4 column. Isocratic elutions were performed with 100 mM phosphate buffer (KH_2_PO_4_), pH 6.5 and 1.2% (v/v) methanol, with a 1 ml/min flow rate. Detection was made at 254 nm. The HPLC equipment used was a Waters® Breeze™ system (Waters Corporation, Milford, Massachusetts, USA). The pump was a Waters® 1525 Binary HPLC Pump, the injector was a Rheodyne™ 7725i (Thermo Scientific, Waltham, Massachusetts, USA) and the detector was a Waters® 2487 Dual Wavelength Absorbance Detector. Software used for data analysis was the Waters® Breeze™ HPLC software. Nucleotide concentrations were determined by interpolation into standard curves for ATP, ADP and AMP. Energy charge was calculated using the following equation ([ATP]+½[ATP])/([ATP]+[ADP]+[AMP]^18^.

Protein quantification was performed using the Bradford method^19^. Bio-Rad® Protein Assay Dye Reagent Concentrate was used with bovine serum albumin (BSA) as standard. After 15 min of incubation in the dark, at 37°C, protein levels were measured by following the absorbance at 595 nm, using BioTek® Cytation3™ UV–vis multi-well plate imaging reader (BioTek, Winooski, Vermont, USA).

#### Assessment of sensitivity to mitochondrial inhibitors

For comparisons of mitochondrial inhibitors’ metabolic effects, intracellular ATP levels were measured using CellTiter-Glo® Luminescent Cell Viability Assay Kit (Promega, Madison, WI, USA), at room temperature and according to the vendor’s recommendations. NHDF cells were plated after trypsinization with cellular densities of 7500 cells/well in 150 μl of culture medium in white 96-well plates. Twenty-four hours after plating, drugs were added to the cells and 3 h later the culture medium was removed, and kit solution was added to the cells. Plates were then subjected to orbital shaking for 2 min, incubated for 10 min, and the luminescent signal was recorded using a BioTek® Cytation3™ UV–vis multi-well plate imaging reader (BioTek, Winooski, Vermont, USA).

#### Analysis of gene expression by quantitative real-time PCR

Total RNA was extracted with PureZOL Reagent, and its concentration and purity were evaluated using a Nanodrop 2000 (ThermoScientific, Waltham, MA, USA). RNA integrity was verified using the Experion RNA StdSens kit (Bio-Rad), and RNA was converted into cDNA using the iScript cDNA synthesis kit (Bio-Rad), following the manufacturer’s instructions. RT-PCR was performed using the SsoFast EvaGreen Supermix, in a CFX96 real time-PCR system (Bio-Rad), with the primers defined in Table 1, at 500 nM. Amplification of 25 ng cDNA was performed with an initial cycle of 30 s at 95 °C, followed by 40 cycles of 5 s at 95 °C plus 5 s at 60°C. At the end of each cycle, EvaGreen fluorescence was recorded to enable determination of Cq. After amplification, melting temperatures of the PCR products were determined by performing melting curves. For each set of primers, amplification efficiency was assessed. Relative normalized expression was determined by the CFX96 Manager software (v. 3.0; Bio-Rad), using the geometric mean of 18S RNA, B2M and HPRT1 as reference. These reference genes were found to be stable between samples from different media. RNA integrity was also assessed using BioRad® PrimePCR™ RNA Control Kit. RNA of a sample was considered intact if |(RQ2 Cq)-(RQ1 Cq)| value (RQ ΔCq) was lower than 3.0. RNA of two samples was considered in a similar degree of integrity if the difference between their RQ ΔCq was lower than 1.0.

**Table 1:**
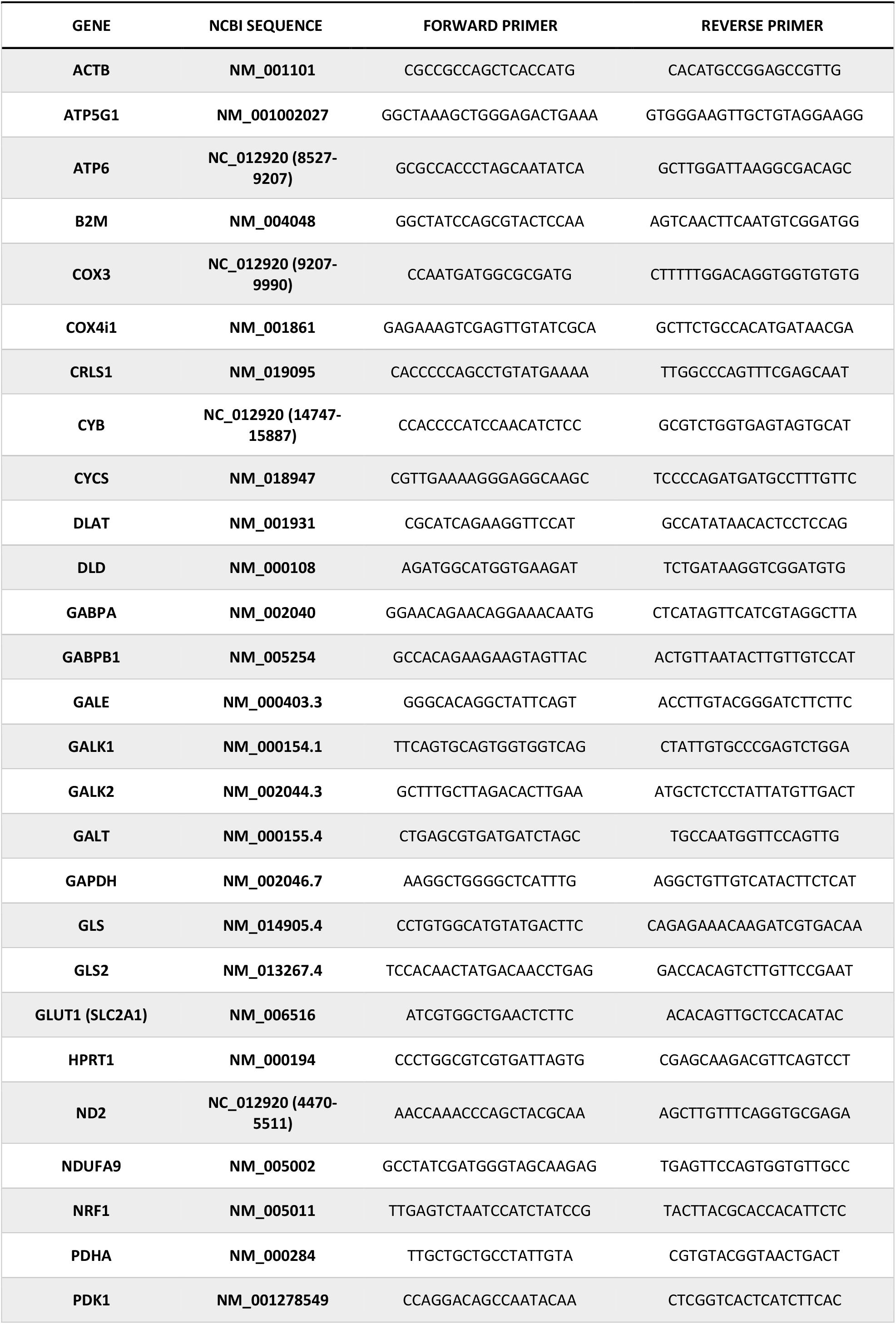
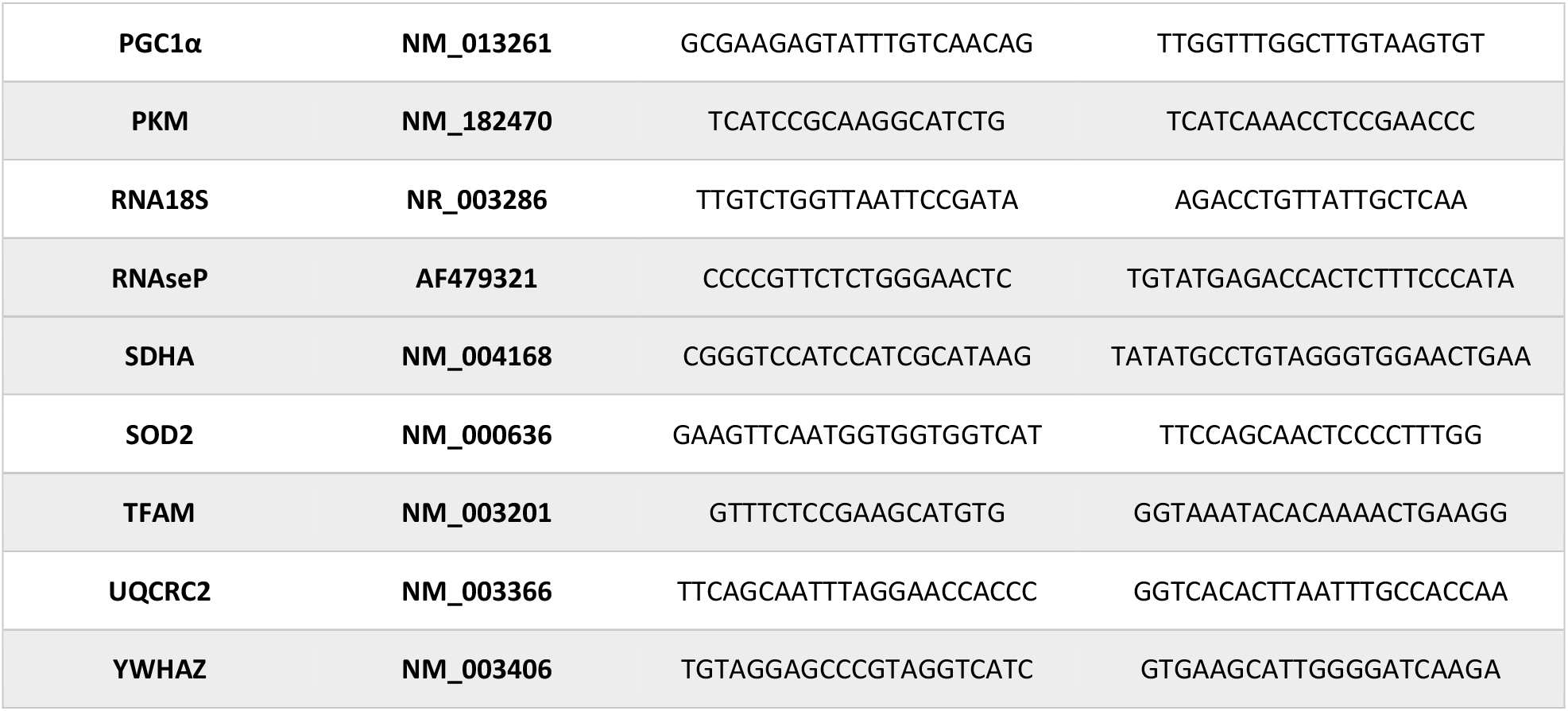
Primer sequence list.

### Data Analysis

Data analysis was performed with Python 3, version 3.7.3, using the Pandas package^20^ to load and transform the data. Information gain between each measured parameter and the target was estimated using the mutual information measure from the Scikit Learn package^21^. Correlation matrices and cluster maps plots were created using the Seaborn module^22^. The correlation between pairs of variables was estimated with the Pearson coefficient. Values span from a total positive (+1) to a total negative (−1) linear correlation, with 0 indicating the lack of linear correlation. Cluster maps were built to group measured parameters obtained for the same sample, and colored to represent the normalized level of expression of a parameter in a specific sample. Parameters and samples were grouped by similarity using a two-way hierarchical clustering method and both dendrograms were drawn using the squared Euclidean distance metric.

Radar plots were generated using Matlab R2020b AddOn spider_plot version 9.0, considering the average of each feature compared to all the samples represented.

Statistical analysis was performed using GraphPad Prism 8.02 software (GraphPad Software, Inc., San Diego, California, USA). Data are represented in bar graphs with dot plots for the number of experiments indicated in figure legends. Statistical significance was set at P<0.05 and determined by Kruskal Wallis method and Dunn’s multiple comparisons test.

## Results

In this work, we tested the mitochondrial and metabolic effects of adapting human adult dermal fibroblasts routinely cultured in high-glucose (HGm) DMEM to either a low-glucose (LGm), or a glucose-free galactose-containing (OXPHOSm) DMEM. This experimental strategy enables us to determine if the absence of glucose induces mitochondrial remodeling and mitochondrial reliance for ATP production, by comparing cells cultured in OXPHOSm with cells cultured in HGm. It also allows us to assess if the reduction of glucose availability to physiological levels is enough to promote mitochondrial remodeling and activity, by comparing LGm with HGm, and if the removal and the reduction of glucose have comparable mitochondrial effects, by comparing OXPHOSm with LGm.

Only OXPHOSm had profound effects on NHDFs bioenergetics, including extensive effects on OCR (**Figure 1**), which were consistent with increased reliance on OXPHOS and reduced reliance on glucose fermentation to meet ATP synthesis demand. In OXPHOSm, cells had substantially increased spare respiratory capacity, basal, ATP-linked, maximal, and non-mitochondrial respiratory rates (**Figure 1D,E**), and only in OXPHOSm, mitochondrial oxidative phosphorylation inhibitors were able to affect intracellular ATP levels (**Figure 1G**). This can result from the fact that cells can overcome respiratory inhibition in glucose-containing media and thus maintain ATP levels by increasing reliance on glycolysis, followed by lactate production. This adaptation is not effective in OXPHOSm, which lacks glucose. Indeed, NHDFs grown in OXPHOSm were far more sensitive to all three respiratory poisons, with significant reductions in ATP levels at concentrations of rotenone, antimycin A, and oligomycin that had no effect in either HGm or LGm. LGm cells showed interesting increases in ECAR, particularly at time points #5-6 (after oligomycin), #9 (after FCCP) and #10-12 (after rotenone + antimycin A) (**Figure 1B,F**).

**Figure 1:**
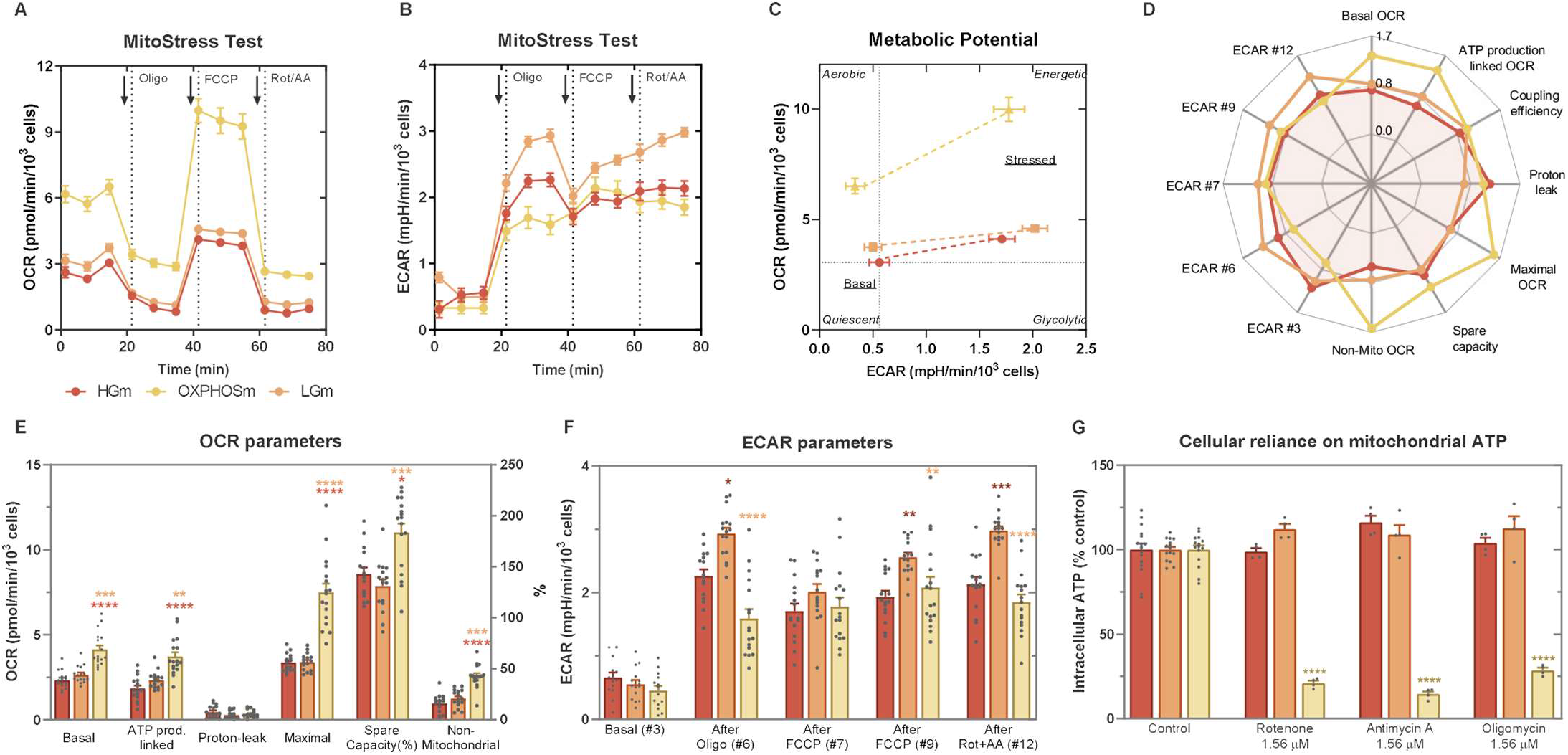
Mitochondrial respirometry and susceptibility to mitochondrial inhibitors in NHDF cells cultured in HGm and adapted to LGm or OXPHOSm. NHDF cells adapted to OXPHOSm, LGm or HGm were plated at a density of 7500 cells/well. **A)** OCR and **B)** ECAR were assessed over time, before (time points #1-3) or after addition of 1) 2 μM oligomycin (time points #4-6), 2) 1 μM FCCP (time points #7-9) and 3) 1 μM antimycin A plus 1 μM rotenone (time points #10-12) and **D-F)** different metabolic parameters were assessed. **C)** Energy map showing metabolic potential of cells before (time point #3) and after being stressed with oligomycin plus FCCP (time point #7). Data represent 3 biological replicates with 4-6 technical replicates. ****: p<0.0001, ***: p<0.001, **: p<0.01, ns: p≥0.05, using Kruskal-Wallis test with Dunn’s multiple comparisons test. **D)** Radar plot summarizing the values obtained for each parameter as a fraction of the average of all measures obtained for the same parameter. **G)** Impact of mitochondrial inhibitors on intracellular ATP content. ATP levels were assessed in cells cultured in HGm or adapted to LGm or OXPHOSm after 3 h of exposure to rotenone, antimycin A or oligomycin. NHDF cells were plated at a density of 7500 cells/well. After 24 h, inhibitors were added to the cells and, 3 h later, ATP levels were measured using CellTiter-Glo® Luminescent Cell Viability Assay Kit. Data are expressed as mean ± SEM of 4 biological replicates. ****: p<0.0001, ***: p<0.001, **: p<0.01, *: p<0.05, compared to the respective control, using ordinary two-way ANOVA test with Tukey’s multiple comparisons test.

Up-regulated mitochondrial bioenergetics promoted by OXPHOSm was accompanied by increased cellular capacity to oxidize TCA cycle substrates, fatty acids and other substrates, including lactate, D-L-a-glycerol phosphate, D-glucose-1-phosphate and D-L-b-hydroxybutyric acid (**Figure 2**). LGm cells seemed to be metabolically similar to HGm cells (**Figure 2B,I**), while OXPHOSm cells manifested a clear metabolic reconfiguration compared to HGm (**Figure 2A**) and LGm (**Figure 2B**), and their substrate-preference profile was not always proportional to that of HGm cells (**Figure 2I**).

**Figure 2:**
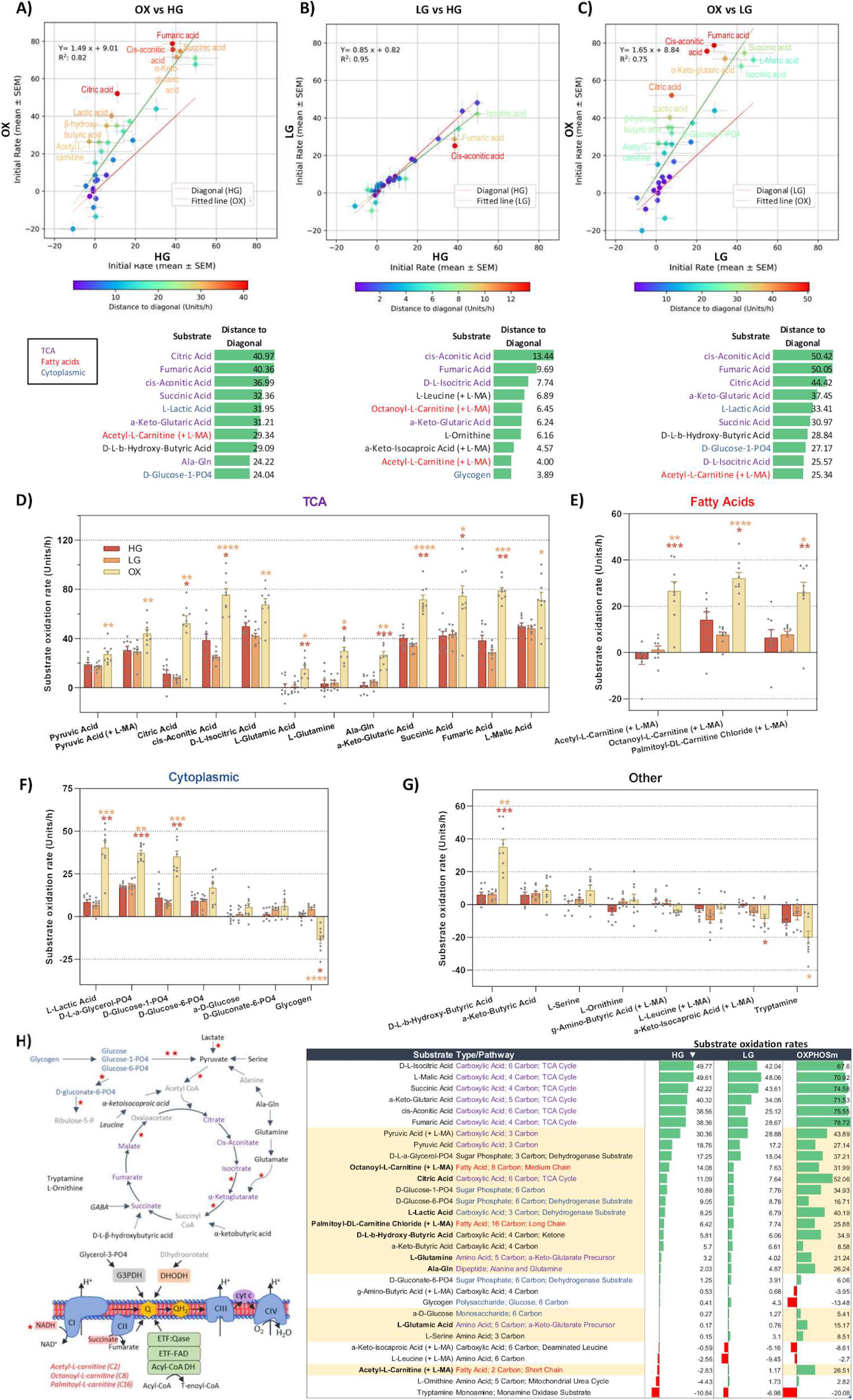
Mitochondrial metabolism and substrate preference. NHDF cells adapted to OXPHOSm, LGm or HGm were plated at a density of 30000 cells/well, and the rate of oxidation of a panel of different types of substrates was assessed using Biolog Mitoplate S1 assay. The rate of oxidation of each substrate was determined and the data were represented in scatterplots to evidence the differences between **A)** OXPHOSm and HGm, **B)** LGm and HGm, and **C)** OXPHOSm and LGm. The color scale represents the difference between the rates observed in the two media for the same substrate, and the top ten substrates and the respective absolute difference are also listed, color coded for the respective pathways **(H)**. Bar charts with dot plots are also shown to evidence the statistical differences between **D)** TCA, **E)** fatty acids, **F)** cytoplasmic and **G)** other substrates. Data represent the mean +-SEM of 3 biological replicates performed in triplicate. ****: p<0.0001, ***: p<0.001, **: p<0.01, ns: p≥0.05, compared to the condition coded with the respective color, using Kruskal-Wallis test with Dunn’s multiple comparisons test. To evidence the changes in substrate preference in the different media, in **I)** substrate oxidation rates were sorted according to the preference of cells cultured in the reference medium, HGm. Substrates for which cell preference changed non-proportionally in OXPHOSm are highlighted in yellow.

One pressing concern we had regarding our approach was that the metabolic stress of acclimation to different media might compromise normal cell physiology. However, we found no evidence that this occurred. Mitochondrial dysfunction can manifest as network fragmentation and/or as increased proton leak with a concomitant decrease in membrane potential and ATP synthesis. However, there was no evidence of network fragmentation in OXPHOSm as mitochondrial networks were generally larger and with longer branches. Similarly, as indirectly inferred from TMRM fluorescence intensity, mitochondrial membrane potential was higher in OXPHOSm than in HGm (though it was slightly lower compared to LGm; **Figure 3F**). Respiration associated with proton leak was not different in cells growing in OXPHOSm compared to other media (**Figure 1 D,E**), and indeed, as a proportion of either basal or maximal OCR, proton leak was lower in OXPHOSm. Similarly, there was no effect on ATP/ADP ratio or cellular energy charge in cells growing in OXPHOSm (**Figure 3 G,H**). Interestingly, the increase in non-mitochondrial OCR observed in OXPHOSm (**Figure 1 D,E**) may be related to an increased pro-oxidative environment, as indicated by increased CM-H_2_DCFDA oxidation in those cells (**Figure 3I**). Overall, our data suggest that NHDFs acclimated to OXPHOSm using the protocol described showed no signs of metabolic stress.

**Figure 3:**
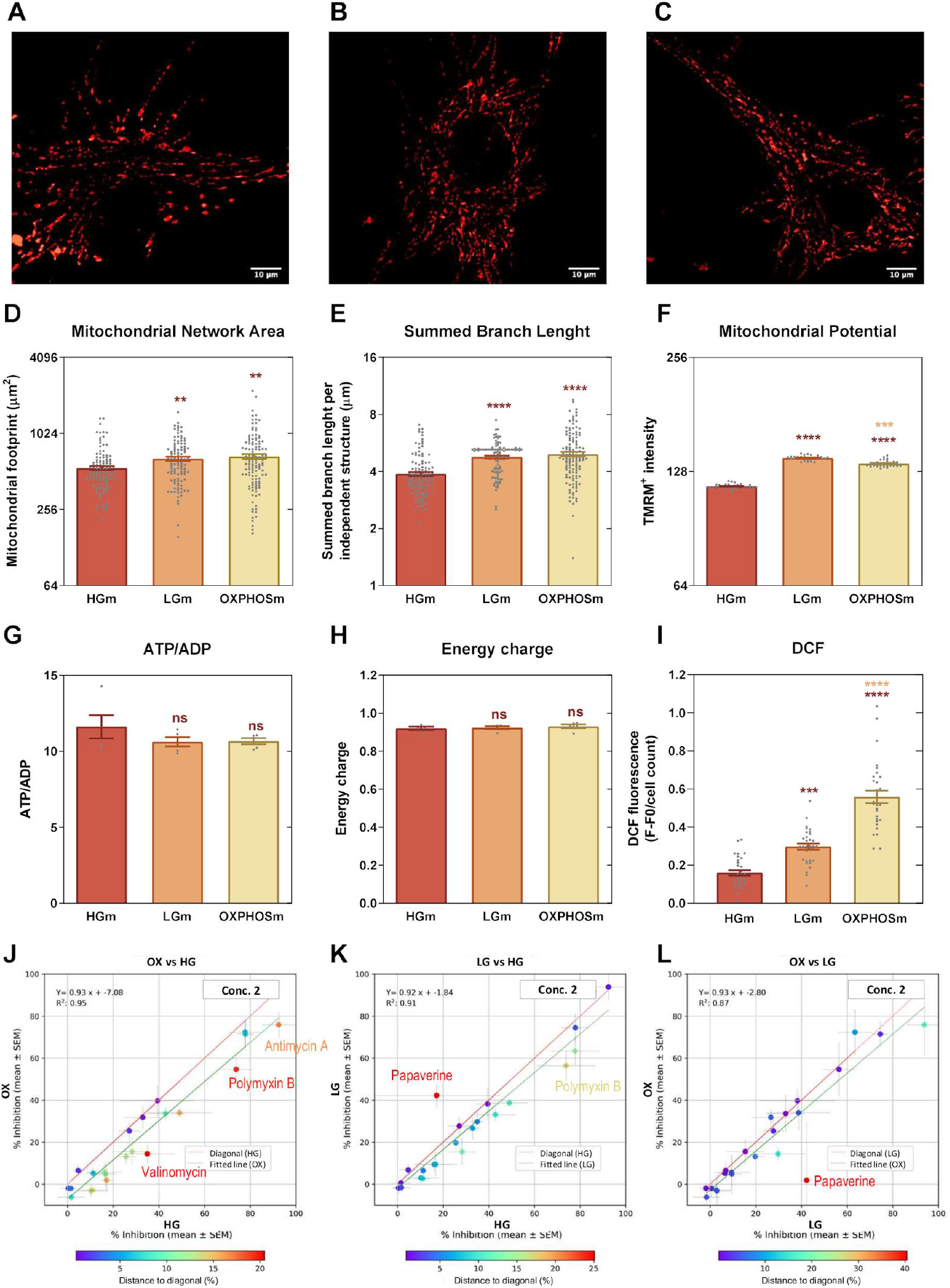
Mitochondrial energy levels and network organization in NHDF cells cultured in HGm and adapted to OXPHOSm or to LGm. Representative images of cells cultured in **A)** HGm, **B)** LGm and **C)**OXPHOSm, labeled with MitoTracker Red, and imaged by confocal microscopy, and then subjected to mitochondrial network analysis using MiNA. **D)** Mitochondrial network area. **E)** Summed Branch Length. **F)** Mitochondrial membrane potential, assessed by measuring TMRM fluorescence intensity. NHDF cells were plated at a density of 2000 cells/well and incubated with 100 nM TMRM and 0.5 μg/ml Hoechst. Imaging was performed by fluorescence microscopy, with 10x amplification. Data represent 3 biological replicates with 8-16 technical replicates. ****: p<0.0001, ns: p≥0.05, compared to the condition with the same color code, using Kruskal-Wallis test with Dunn’s multiple comparisons test. **G)** ATP/ADP ratio and **H)** Energy Charge in NHDF cells cultured in HGm or adapted to LGm or OXPHOSm. ATP, ADP and AMP were measured by HPLC. Data represent 4-5 independent experiments. ns: p≥0.05, using Kruskal-Wallis test with Dunn’s multiple comparisons test. **I)** Analysis of redox environment was assessed by following the oxidation of CM-H_2_DCFDA by fluorimetry. Data represent 4-5 independent experiments and are expressed as mean ± SEM. ****: p<0.0001, ***: p<0.001, compared to the condition with the same color code, using Kruskal-Wallis test with Dunn’s multiple comparisons test. **J-L)** Inhibition of mitochondrial succinate oxidation by a panel of different mitochondrial inhibitors was assessed using Mitoplate I1 assay. Scatterplots represent the mean +-SEM of the rate of inhibition in the presence of each inhibitor at the second highest concentration. The color scale represents the difference between the rates observed in the two media for the same substrate.

Curiously, despite no substantial changes in OCR parameters, the mitochondrial network area and branch length were increased in LGm, which may also explain the higher intensity of TMRM fluorescence already described (**Figure 3 D,E**). This seems to indicate that LGm promotes an increase in mitochondrial mass, but not in mitochondrial activity. To further understand this, we used the MitoPlate I-1 assay (**Figure 3 J-L** and **Supp Figure 3**), to study the effects of four increasing concentrations of different mitochondrial inhibitors on the rate of mitochondrial succinate oxidation, for a fixed number of saponin-permeabilized cells previously cultured in the three different media. Since cells were permeabilized, the maximum degree of succinate oxidation in cells cultured in each medium depended mainly on the abundance and activity of mitochondrial machinery. A higher inhibition of succinate oxidation for the same concentration of the inhibitor is indicative of lower mitochondrial abundance or activity of components downstream succinate oxidation. A differential response to specific inhibitors may also inform on the presence or activity of specific metabolic pathways. Considering for example the second concentration of inhibitors, for which a better discrimination was obtained, lower inhibition was observed when comparing LGm to HGm or OXPHOSm to HGm, whereas LGm and OXPHOSm were more similar (**Figure 3J-L** and **Supp Figure 3**). This is in agreement with the observed increase in mitochondrial network parameters in both LGm and OXPHOSm. Regarding specific inhibitors, a curious response to papaverine was observed in LGm cells, in which mitochondrial succinate oxidation was less affected by this compound.

Furthermore, we wanted to understand how transcriptional changes shaped the metabolic and morphological mitochondrial remodeling. Gene expression analysis was performed by qPCR using a panel of 33 genes of interest belonging to different functional categories (OXPHOS subunits, Mitochondrial Biogenesis, Metabolic Enzymes, Galactose Metabolism and General Cell Functions) and 3 reference genes (RNA18S, B2m and HPRT1) (**Figure 4**).

**Figure 4:**
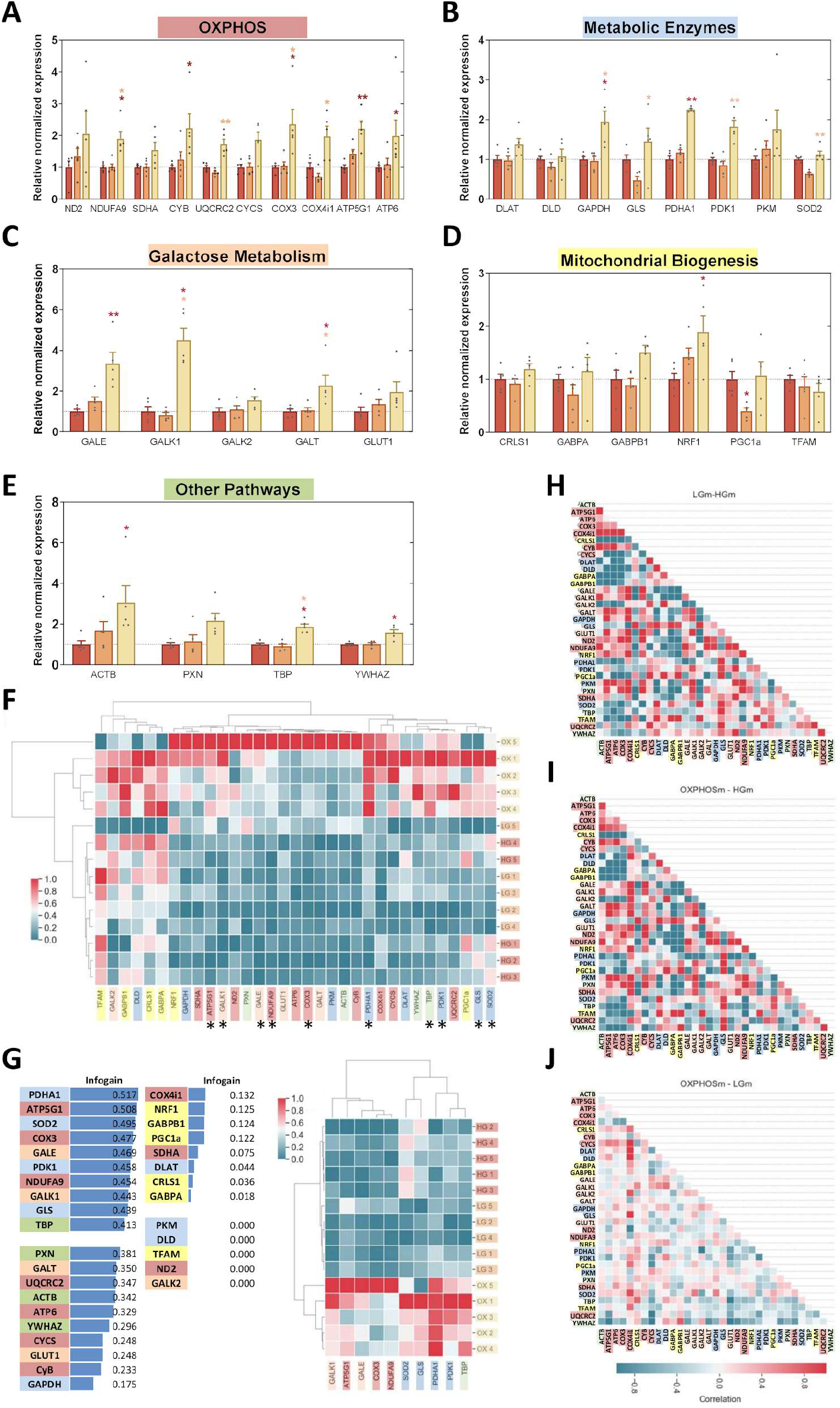
Changes in gene expression in cells cultured in HGm or adapted to LGm or OXPHOSm. Total RNA was extracted, purified, converted into cDNA, and amplified by qRT-PCR. Gene expression was measured and normalized to the geometric mean of B2M, HPRT1 and RNA18S levels, and divided by the mean of the values obtained for HGm-cultured cells. Genes were grouped in five different categories, **A)** OXPHOS subunits, **B)** Metabolic Enzymes, **C)** Galactose Metabolism, **D)** Mitochondrial Biogenesis and **E)** Other Pathways. Data represent 5 biological replicates. **: p<0.01, *: p<0.05, compared to the condition with the same color code, using Kruskal-Wallis test with Dunn’s multiple comparisons test. **F)** Data are presented as clustergrams of gene expression by target gene and sample. Relative expression levels are color coded from white (no regulation), blue (downregulation) to red (upregulation). Darker color shades represent greater relative expression differences. **G)** Importance of each gene for identification of cell culture medium was determined by calculating the Information gain (Infogain). The 10 more informative genes are labeled with a star in the clustergram. Differential correlation matrices for gene expression values obtained in cells cultured in **H)** LGm compared to HGm, **I)** OXPHOSm compared to HGm and **J)** OXPHOSm compared to LGm. Correlation differences are represented in a color scale. The respective individual correlation matrices are represented in **Supplemental Figure 2**.

Increased expression of mitochondrial OXPHOS complexes I, III, IV, and V subunits and PDHA1 was observed in OXPHOSm, consistent with increased mitochondrial mass (**Figure 4A,B**), although some of the major players in mitochondrial biogenesis remained unchanged (**Figure 4D**). However, an increase in NRF1 transcripts was observed in OXPHOSm cells (**Figure 4D**), possibly mediating the observed increase in mitochondrial mass. The expression of GALK1, GALT, and GALE increased significantly (**Figure 4C**) in OXPHOSm, suggesting a substantial induction of the Leloir pathway for galactose catabolism. Increased transcripts for genes commonly used as reference genes in the literature were also observed in OXPHOSm cells (**Figure 4E**).

Data were integrated considering the levels of all transcripts for each independent sample. For gene expression data, the transcripts with higher information gain regarding the culture medium were PDHA1, ATP5G1, SOD2, COX3, GALE, PDK1, NDUFA9, GALK1, GLS and TBP (**Figure 4G**). Using transcriptional data, we analyzed the correlations between transcripts in each target group (Supplemental Figure 2) and calculated the correlation differences between LGm and HGm, OXPHOSm and HGm and OXPHOSm and LGm (**Figure 4H-J**). LGm and OXPHOSm presented significant differences in transcripts’ correlations compared to HGm, whereas OXPHOSm and LGm seemed to be more similar, with only a few differences observed. This fact contrasts with the lack of changes regarding bioenergetic parameters, but is consistent with a similar remodeling of mitochondrial content and morphology.

A clustering analysis using only the 10 most informative genes previously identified provided a complete distinction between samples from the 3 culture media (**Figure 4G**). The genes formed two clusters, one containing only genes related to OXPHOS subunits and galactose metabolism, and another containing 4 metabolic enzymes and TBP. This analysis revealed that gene expression in LGm- and OXPHOSm-cultured cells followed a completely different transcriptional pattern compared to cells cultured in HGm, and allowed us to identify a subset of transcripts that are characteristic of cells growing in each of the 3 different media. These results indicate that a short protocol, in which gradual acclimation to OXPHOSm was made during the first passage, was sufficient to ‘re-wire’ NHDF metabolism toward a more oxidative, mostly glucose-independent, metabolism. To understand which components of OXPHOSm were important for establishing the respiratory phenotype, we systematically removed key constituents. Removal of galactose had little effect on OCR parameters (**Figure 5**). Neither basal, ATP-linked, or maximal respiration rates, nor spare respiratory capacity of NHDFs were different between OXPHOSm and OXPHOSm lacking galactose. Both ECAR and stressed ECAR were lower in OXPHOSm-Gal, suggesting that galactose-derived glucose-1-phosphate is metabolized until conversion of pyruvate to lactic acid. In contrast to galactose, removal of glutamine strongly reduced most OCR values, indicating that it is indispensable to the oxidative phenotype in these cells. Removal of pyruvate alone was of no consequence to OCR or ECAR, while simultaneous removal of both galactose and pyruvate increased basal respiration and proton leak, but decreased maximal and non-mitochondrial respirations, as well as spare capacity (**Figure 5**).

**Figure 5:**
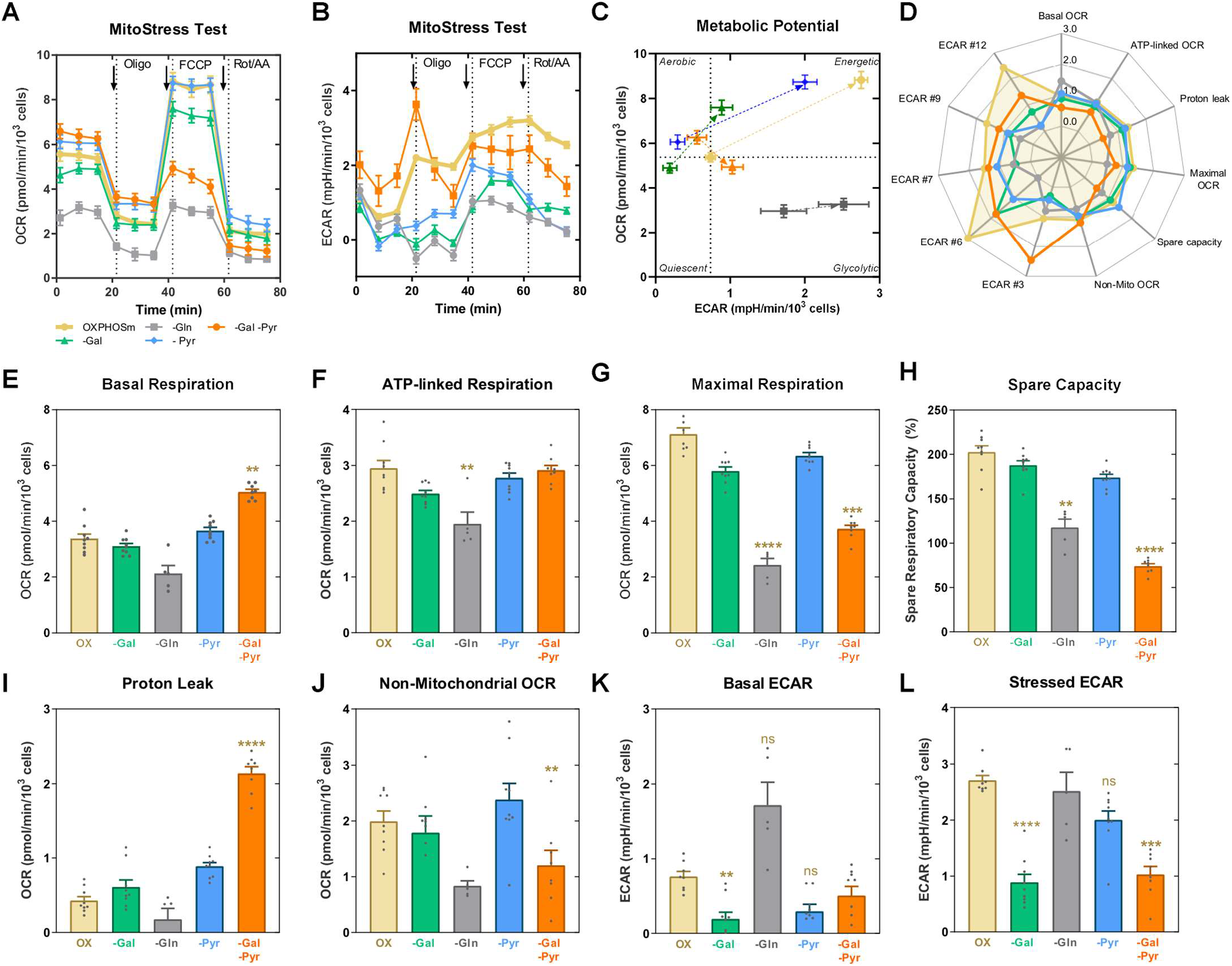
Mitochondrial Stress Test featuring OCR- and ECAR-associated parameters in NHDF cells cultured in OXPHOSm in the presence or in the absence of galactose, glutamine, pyruvate and glutamine plus pyruvate. NHDF cells were plated at a density of 7500 cells/well. **A)** OCR and **B)** ECAR of NHDF cells adapted to OXPHOSm or its variations was assessed over time, before or after addition of 1) 2 μM oligomycin, 2) 1 μM FCCP and 3) 1 μM antimycin A plus 1 μM rotenone and **E-L)** different metabolic parameters were assessed. **C)** Energy map showing metabolic potential of cells when stressed with oligomycin plus FCCP. **D)** Radar plot summarizing the values obtained for each parameter as a fraction of the average of all measures obtained for the same parameter. Data represent 5-9 replicates. ****: p<0.0001, ***: p<0.001, **: p<0.01, ns: p≥0.05, using Kruskal-Wallis test with Dunn’s multiple comparisons test.

## Discussion

Cell culture conditions enormously influence metabolism *in vitro*, which is highly relevant for preclinical assays. Fibroblasts are an interesting cell model for preclinical applications^2, 12^, namely regenerative medicine, diagnostics and therapeutic development for personalized medicine as well as in the validation of ingredients for cosmetics.

Stimulation of OXPHOS in cultured cells can unmask specific mitochondrial toxicants and has gained increased interest for the study of mitochondrial health in Toxicology and Biomedicine. Unmasking selective mitochondrial toxicity is crucial for toxicological studies and the correct identification of mitochondrial toxicants is known to be influenced by cell culture conditions^7,23, 24, 25^.

Here, we aimed to optimize a protocol for mitochondrial health studies in human skin fibroblasts that is compatible with their short lifespan, since previous studies used adaptation protocols requiring most of the replicative lifespan of the cells^12^, shortening the window of opportunity for experiments. Thus, we wanted to reduce as much as possible the period of cellular adaptation to media of different composition, although not compromising the detection of mitochondrial toxicity was promoted. Subsequently, we characterized metabolic remodeling in cells that are more susceptible to mitochondrial toxicants, to find biomarkers of mitochondrial health, and understand the importance of the available energy substrates for mitochondrial morphological and metabolic remodeling. We assessed whether it was enough to reduce glucose availability to promote OXPHOS and uncover the effects of mitochondrial inhibition, and what was the role of other energy substrates in the culture medium.

### OXPHOSm vs HGm: Removal of glucose and replacement with galactose induces mitochondrial remodeling and cellular reliance on mitochondria for ATP production

Glucose-free media have been previously used to improve the correct identification of mitochondrial inhibitors, such as complex I inhibitor rotenone, complex III inhibitor antimycin A and complex V inhibitor oligomycin^7, 12, 26^. Here, we used a short term adaptation protocol to glucose-free OXPHOSm, which successfully uncovered the mitochondrial effects of these three OXPHOS inhibitors in NHDF cells.

To understand the metabolic remodeling underlying increased mitochondrial stress in OXPHOSm-cultured NHDF cells, we analyzed OCR- and ECAR- associated parameters (**Figure 1**) and mitochondrial substrate preference (**Figure 2**). Most of OCR-associated parameters, namely Basal, ATP production-linked, Maximal and Non-Mitochondrial respirations significantly increased in OXPHOSm-adapted cells compared to with HGm-cultured cells, demonstrating that cells adapted to OXPHOSm were more oxidative, as expected. Under mitochondrial stress promoted by oligomycin and FCCP (time point #7), OXPHOSm-adapted cells became much more energetic (less quiescent) than the others, increasing both ECAR and OCR compared to basal conditions (time point #3)(**Figure 1C**). The increase in OCR parameters was associated with a general increase in the cellular capacity to oxidize TCA cycle substrates, but also with an increased preference for fatty acids, lactate and glycerol phosphate (**Figure 2**). Additionally, the enhanced Non-Mitochondrial respiration suggests that the metabolic remodelling also occurs in other organelles, for example due to increased generation of reactive oxygen species (ROS). A possibility is that not only mitochondrial but also the peroxisomal β-oxidation is stimulated, since there is an interplay between both metabolic pathways^27^. Peroxisomal β-oxidation is dependent on O_2_ consumption, which is reduced to H_2_O_2_^27, 28^. Thus, if this assumption is true, the peroxisome-derived H_2_O_2_ can also explain the increased pro-oxidant environment in OXPHOSm-adapted, more metabolically active, cells (**Figure 3I**). Further work to confirm this hypothesis is required.

The next step was to evaluate whether metabolic remodeling of NHDF cells after adaptation to OXPHOSm affected general cellular energetic state, by determining ATP/ADP ratio and Energy Charge, which may inform on the cellular energy levels^18, 29, 30, 31^. ATP/ADP ratio and energy charge values were similar in all the conditions tested (**Figure 3**), indicating that, regardless of the substrates available for metabolization, cells adapted to use them to produce ATP, maintaining the energetic state independently of those sources.

The availability of different energy substrates may impact mitochondrial network structure, which in turn affects the efficiency of ATP production^32^. Thus, we wanted to understand whether the changes in mitochondrial function in OXPHOSm-adapted cells were associated with changes in mitochondrial area, morphology and/or polarization. A higher mitochondrial area per cell was observed after adaptation to OXPHOSm than in cells cultured in HGm (**Figure 3**). These results were the opposite to what was observed in HeLa cells, in which cellular area was higher in cells grown with glucose. In the same work, mitochondrial content was also not increased in OXPHOSm-adapted cells, and mitochondria in OXPHOSm-adapted cells were more elongated^33^, which was also observed in our group using BJ fibroblasts^12^. Furthermore, mitochondrial membrane polarization, estimated by TMRM intensity^34^, was increased in OXPHOSm-adapted NHDFs (**Figure 3B**).

Previous studies described increased gene expression of several mtDNA-encoded OXPHOS subunits after adaptation to OXPHOSm^12^ and also in protein levels of some subunits^12, 33^. Our results show that adaptation to OXPHOSm promoted significant increases in the expression of the mitochondrial-encoded CYB (complex III), COX3 (complex IV) and ATP6 (complex V) genes (**Figure 4**), as well as in nuclear DNA-encoded NDUFA9 (complex I) and ATP5G1 (complex V) genes (**Figure 4**), when compared to the expression of these genes in HGm-grown cells.

The expression of several metabolism-related transcripts was also affected. GALK1, GALE and GALT were upregulated in cells adapted to OXPHOSm, suggesting that cells adapted to metabolize galactose in the absence of glucose. This is consistent with what was observed in BJ cells cultured in OXPHOSm, which also showed increases in GALE and GALK1 expression^12^. Interestingly we also observed a significant increase in GAPDH gene expression, as well as a tendency for increased PKM expression (**Figure 4**), both related to glycolytic metabolism. Accordingly, OXPHOSm-adapted cells were also more capable to oxidize glucose-1-phosphate than HGm cells (**Figure 2**). Although this may seem unexpected, the glycolytic system was previously shown to be similar in HeLa cells grown with or without glucose^33^. This is also supported by the small, non-significant, increase in gene expression of GLUT1 transporter in OXPHOSm-adapted cells (**Figure 4**), responsible not only for glucose but also for galactose uptake^35, 36^. Given that GLUTs have generally lower affinity for galactose than for glucose^37^, an increase in GLUT1 transcripts may be associated with cellular adaptations to achieve increased glycolytic efficiency.

We also evaluated genes associated with the metabolism of the other energy substrates present in the culture medium, namely, glutamine and pyruvate. No differences in GLS gene expression in OXPHOSm-adapted and in HGm-cultured cells, although a non-statistically significant increase was observed (**Figure 4**). Regarding the PDH complex, PDHA1 transcript levels were significantly increased (**Figure 4**), suggesting an increase in pyruvate channeling towards the TCA cycle. An increase in PDH protein levels was previously observed in cells grown without glucose^33^. Interestingly, gene expression of PDK1, a negative regulator of PDH and a member of the PDH complex, showed a non statistically significant increase (**Figure 4**). This increase accompanied PDHA1 increased expression and could mean a compensatory increase to regulate PDH activity.

Together with the increased gene expression of many OXPHOS subunits in OXPHOSm samples, the results suggest that mitochondrial biogenesis occurred in OXPHOSm-adapted cells.

### LGm vs HGm: Reduction of glucose availability to physiological levels is enough to promote mitochondrial network remodeling but not metabolic activity

Although the reduction of glucose availability (LGm) did not affect OCR parameters nor mitochondrial substrate preference, it increased ECAR after metabolic challenges with oligomycin, FCCP and rotenone plus antimycin A (**Figure 1**). LGm also seemed to promote a morphological remodeling of the mitochondrial network, with increased mitochondrial transmembrane potential. Inhibition of succinate oxidation by a panel of compounds with different mechanisms of action was less effective in LGm cells, also suggesting higher mitochondrial content. An imbalance of the redox equilibrium towards a more oxidative environment was also observed in LGm cells, although significantly less expressive than in OXPHOSm-adapted cells.

The reduction of glucose availability by using LGm induced almost no changes in the measured transcript levels compared to HGm, apart from a significant decrease in PGC1a levels (**Figure 4**). This favors the idea that, metabolically, cells cultured in LGm or HGm are very similar. However, although it was not explicit when looking at the levels of the transcripts analyzed, the correlations between levels of pairs of transcripts were highly affected by LGm, in a way that was similar but did not overlap with OXPHOSm-adapted cells.

Our results show that cells cultured in LGm or HGm are very similar metabolically, and thus HGm should not be used in regular screening assays using these cells, as LGm represents a more physiological situation.

### OXPHOSm vs LGm: Removal and reduction of glucose do not have comparable effects

Statistically significant differences between LGm and OXPHOSm-adapted cells were found in Basal, ATP-linked, Maximal and Non-Mitochondrial respirations, and Spare Respiratory Capacity, all higher in OXPHOSm-cultured cells (**Figure 2**).

Differences in the transcriptional profiles of OXPHOSm and LGm were limited to a few gene pairs, particularly involving correlations that include the genes COX4i1, CYCS and YHWAZ (**Figure 4D**). Our results show that the removal and reduction of glucose do not have comparable effects. Culture medium should be carefully chosen to ensure that cells are metabolically capable.

### OXPHOSm vs LGm vs HGm

Considering all the gene expression data, the most informative genes to distinguish between samples from the 3 culture media were PDHA1, ATP5G1, SOD2, COX3, GALE, PDK1, NDUFA9, GALK1, GLS and TBP (**Figure 4**). A clustering analysis using only these 10 most informative genes provided a complete distinction between samples from the 3 culture media (**Figure 4**), which formed two gene clusters, one containing only genes related to OXPHOS subunits and Galactose metabolism, and another containing 4 metabolic enzymes and TBP. TBP is commonly used as a reference gene for qPCR experiments. Our results show that TBP, and also ACTB and YWHAZ (**Figure 4**), should not be used for normalization of gene expression in samples from cells cultured in OXPHOSm, at least when comparing them with cells cultured in HGm or LGm. Thus, we identified a subset of critical markers that allowed for a perfect distinction between samples obtained in the 3 different media (**Figure 4**), which will help select the best culture media to evidence different types of mitochondrial stresses.

Curiously, genes related to mitochondrial biogenesis were not very informative to distinguish between samples from the 3 media, although a significant increase in NRF1 levels was observed in OXPHOSm samples and decreased PGC1a was observed for LGm (**Figure 4**).

### Culture medium components and mitochondrial activation

Our results clearly show that decreasing glucose availability affects mitochondria, but is not the same as replacing glucose by galactose. However, it is not clear if the presence of galactose is important, or if it is the absence of glucose that plays the central role in promoting mitochondrial function. Thus, we wanted to understand the role played by the other energy substrates present in OXPHOSm, namely glutamine, galactose and pyruvate.

Glutamine is both a cellular building block and an indirect fuel for the TCA cycle^38, 39^ and was described as the primary energy fuel for HeLa cells^40^. Depending on the cellular metabolic state, glutamine can be used as either a biosynthetic precursor for proteins, nucleotides or glutathione, or as an anaplerotic fuel that drives mitochondrial ATP production^41, 42^. Without its presence in glucose-free culture medium, it is difficult to predict the adaptations in energy metabolism of NHDF cells. OCR- and ECAR- associated parameters were measured in NHDF cells adapted to OXPHOSm or to OXPHOSm-Gln (**Figure 5**), whose composition is similar to OXPHOSm but without glutamine. In the absence of glutamine, there were significant decreases in ATP Production-linked, Maximal and Non-Mitochondrial Respirations and in Spare Respiratory Capacity. These results suggest that glutamine is essential for the stimulation of OXPHOS in NHDF cells adapted to glucose-free media, and limits their potential to rely on OXPHOS in stress situations. In fact, in cells adapted to OXPHOSm-Gln, the full ETC capacity was limited, since Spare Respiratory Capacity was only slightly higher than 115%, far from the ~200% observed for cells adapted to OXPHOSm. Basal ECAR showed a tendency to increase in OXPHOSm-Gln -adapted cells, which may reflect the fact that galactose and/or pyruvate might be converted in lactate in a higher extent in these cells, since their oxidative metabolism is repressed due to the absence of glutamine.

Galactose seemed to contribute to OXPHOS induction in NHDF cells adapted to OXPHOSm. Galactose metabolization generates two NADH molecules, which can be oxidized in mitochondria through OXPHOS^43^. In HeLa cells, glycolysis was shown to be dispensable for ATP production in cells grown in glutamine-containing media^40^. Thus, it is interesting to assess the need of sugar in the culture medium to support NHDF cells. For that purpose, OCR- and ECAR- associated parameters were measured in NHDF cells adapted to OXPHOSm or to OXPHOSm-Gal, which lacks galactose. We observed that Basal and Proton Leak-associated Respirations were significantly increased in cells adapted to OXPHOSm -Gal when compared with OXPHOSm-grown cells (**Figure 5**). Results also seem to suggest that galactose is somehow important for OXPHOS promotion in NHDF cells, since ATP Production-Linked and Maximal Respirations of cells adapted to OXPHOSm-Gal were slightly decreased compared to cells adapted to OXPHOSm. This result might appear difficult to explain at first sight since galactose contribution for OXPHOS would be small, given the presence of exogenous pyruvate and glutamine, according to previous reports^40^. However, in the absence of sugars in the culture medium, there is no carbon source for the pentose phosphate pathway (PPP), which is probably an important fate of galactose^40^. Hence, many biosynthetic reactions are compromised without galactose as the carbon source, including those that produce NADH or ATP, which have PPP intermediates as precursors^44^. In some cell types, it is possible to speculate that this absence of sugar could eventually be compensated by stimulating gluconeogenesis. However, this cannot be the case in NHDF cells, since gluconeogenesis key enzyme phosphoenolpyruvate carboxykinase-C (PEPCK-C) was reported not to be expressed in the skin^45^, underlining the importance of the presence of galactose for NHDF cells grown in media without glucose. This may not be the case for cell lines derived from tissues where PEPCK-C is highly expressed^45^. Further metabolomic analyses are required to understand the exact role of galactose for NHDF cell metabolism grown in these conditions.

Pyruvate seemed not to be critical for OXPHOS induction in cells grown without glucose and with galactose supplementation. Pyruvate can be converted into acetyl-CoA and enter the TCA cycle, thus being an energy source for cells^46^. However, if pyruvate is the only source of energy, this will probably only work in short-term because there would be no source to maintain the pool of TCA cycle intermediates that are used in other metabolic pathways, as opposed to what happens when glutamine is the main fuel^41^. Pyruvate supplementation in culture media without glucose is not entirely consensual. Pyruvate is not only a carbon source for the TCA cycle, and therefore for the synthesis of metabolites that feed ETC^46^, but can also act as ROS scavenger^47, 48, 49, 50^, contributing to a lower degradation of the mitochondrial proteins and hence to a possible increased enzymatic activity of the ETC complexes. Nevertheless, supplementation of the culture medium with pyruvate, in addition to a sugar, is often considered redundant, since the metabolism of sugars through glycolysis produces pyruvate^46^. Our data support this hypothesis, that seems to indicate that pyruvate supplementation has negligible influence in OXPHOS induction indeed, since its absence in culture OXPHOSm-Pyr did not lead to a decrease in most OCR-associated parameters, such as basal, ATP production-linked, maximal and non-mitochondrial respirations (**Figure 5**). These results are probably due to metabolic redundancy. Although pyruvate supplementation is unnecessary for OXPHOS stimulation, its removal results in a big decrease in its pool, which can only be maintained by galactose metabolization. Thus, given the decrease and the constant demand of pyruvate for the TCA cycle, the amount of pyruvate metabolized into lactate should be lower in cells cultured in OXPHOSm-Pyr, resulting a medium acidification comparable to the one observed for OXPHOSm.

Galactose and pyruvate seemed to be mainly dispensable for the metabolic remodeling toward OXPHOS. Thus, it is interesting to understand what happens if both of them are removed. OXPHOSm -Gal -Pyr was the simplest medium tested, since it was supplemented only with glutamine. These cells presented the highest Basal Respiration value of all the conditions analyzed, and their ATP production-linked OCR value was similar to the value observed for cells adapted to OXPHOSm (**Figure 5**). These values indicate that, indeed, these are probably the most oxidative cells analyzed. However, these cells did not increase their OCR in response to FCCP, resulting in their Maximal Respiration being lower than their Basal Respiration (**Figure 5**). This behavior may be due to increased sensitivity to oligomycin and FCCP toxicity, resulting in the induction of necrosis. Since these cells relied solely on glutamine for ATP production, the higher susceptibility to stress observed in cells cultured in OXPHOSm -Gal -Pyr may be related to the generation of ammonia as a metabolic waste product during glutamine catabolism, which is potentially toxic^51^.

## Conclusions

In this work, we optimized a protocol for mitochondrial fitness studies in human skin fibroblasts, which is compatible with their short lifespan and uncovers mitochondrial inhibitors’ effects. The metabolic remodeling occurring in cells that are more susceptible to mitochondrial toxicants revealed mitochondrial performance biomarkers, including genes associated with carbohydrate metabolism and oxidative phosphorylation. We also demonstrated the importance of the available energy substrates for OXPHOS stimulation, and found that reducing glucose availability is not enough, whereas glutamine is the primary fuel, as long as galactose and/or pyruvate are also available.

Culturing fibroblasts in OXPHOSm including a short-term adaptation period should be useful to test the safety of drugs intended for the skin, for testing safety and efficacy of cosmetic ingredients, for the identification and correction of mitochondrial defects in fibroblasts obtained from patients and potentially also for the metabolic priming of fibroblasts for regenerative medicine.

## Acknowledgements

This work was financed by the European Regional Development Fund (ERDF), through the COMPETE 2020 - Operational Programme for Competitiveness and Internationalization and Portuguese national funds via FCT – Fundação para a Ciência e a Tecnologia, under projects PTDC/BTM-SAL/29297/2017-POCI-01-0145-FEDER-029297, POCI-01-0145-FEDER-029391-PTDC/MED-FAR/29391/2017, UIDB/04539/2020, DL57/2016/CP1448/CT0016 and SFRH/PD/BD/143055/2018. We are thankful to Barry Bochner and Enrico Tatti from Biolog for making the Omnilog device available for this study and for insightful comments on our data.

