## Supplemental data for "Mitochondrial and metabolic remodeling in human skin fibroblasts in response to glucose availability"

Supplementary Figures

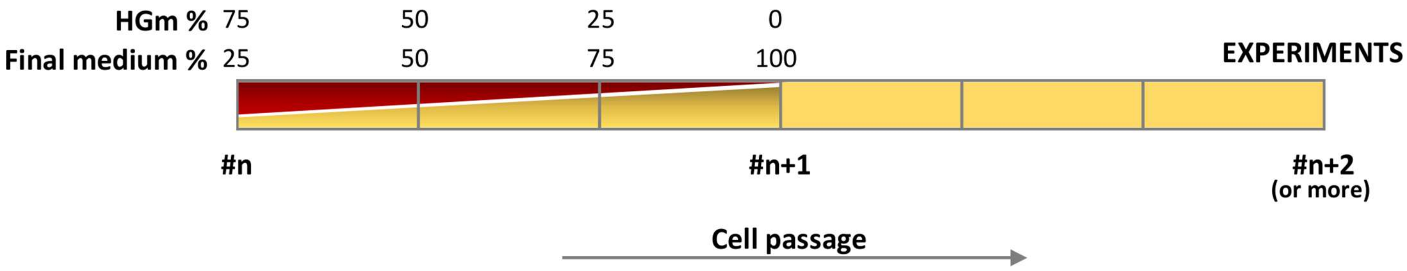


***Supplemental Figure 1: Schematic representation of adaptation protocol used in this study.*** *NHDF cells were routinely cultured in HGm. For the experiments part of the cells were gradually adapted to new culture media during one cell passage and the experiments were carried out in the final medium.*

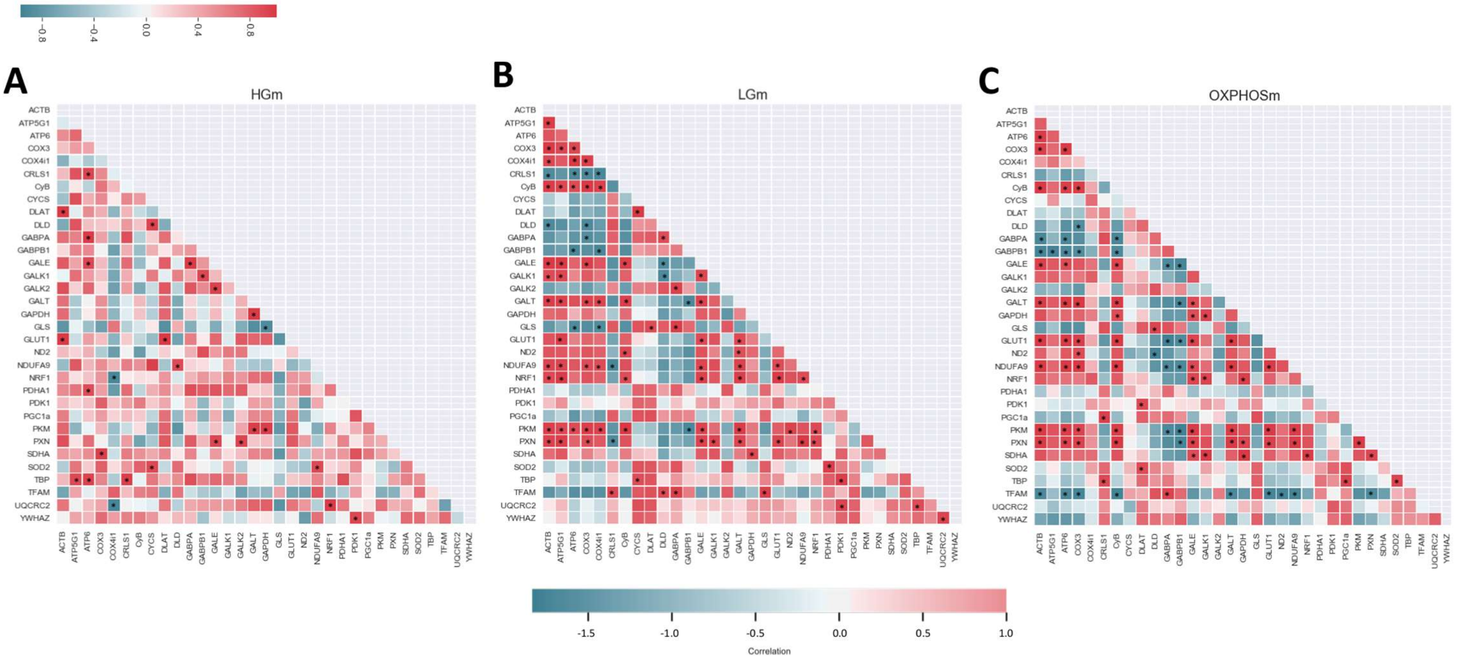


***Supplemental Figure 2: Correlation matrices for gene expression values obtained in cells cultured in A) HGm B) LGm and C) OXPHOSm.*** *Correlation values are represented in a color scale and stars represent correlations with p<0.05. The differential correlation matrices are represented in **Figure 4**.*

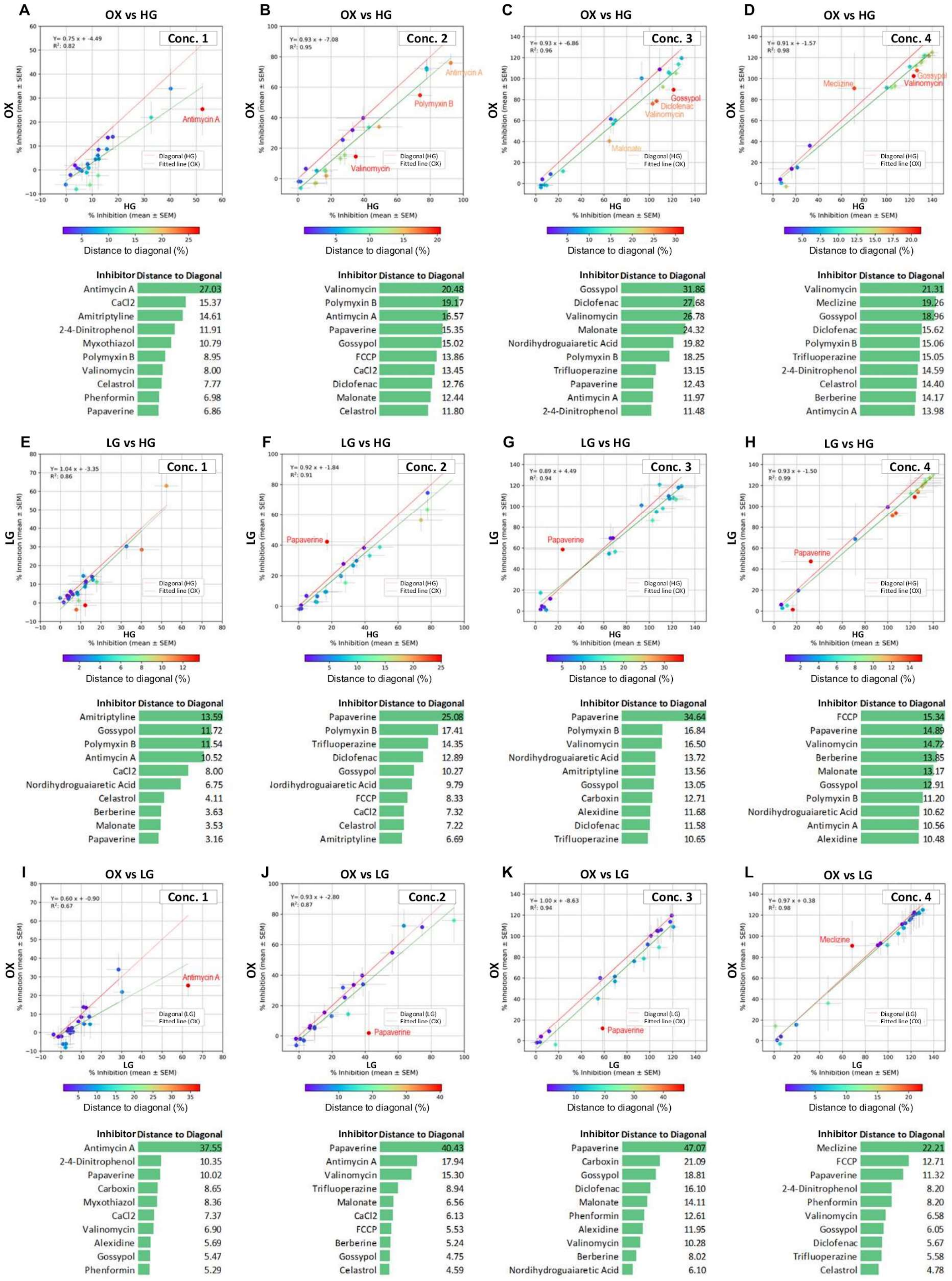


***Supplemental Figure 3: Inhibition of mitochondrial succinate oxidation by a panel of different mitochondrial inhibitors was assessed using the Mitoplate I1 assay.*** *Scatterplots represent the mean +-SEM of the rate of inhibition in the presence of each inhibitor at four increasing concentrations in two different media. The color scale represents the difference between the rates observed in the two media for the same inhibitor. For each graph, the top ten inhibitors with higher distances to diagonal and the respective absolute differences (in % inhibition) are also listed.*
